# Genetic, Natal, and Spatial Drivers of Social Phenotypes in Wild Great Tits

**DOI:** 10.1101/2024.07.11.603055

**Authors:** Devi Satarkar, Irem Sepil, Ben C. Sheldon

## Abstract

In social animals, group dynamics profoundly influence collective behaviours, vital in processes like information sharing and predator vigilance. Disentangling the causes of individual-level variation in social behaviours is crucial for understanding the evolution of sociality. This requires unravelling the genetic and environmental basis of these behaviours, which is challenging in uncontrolled wild populations. In this study, we partitioned genetic, developmental and spatial environmental variation in repeatable social network traits derived from foraging events using a multigenerational pedigree and extensive observational social data from a long-term monitored great tit population. Animal models indicated minimal narrow-sense heritability (2-3%) in group size choice, further reduced when spatial location was considered, which itself explains a substantial 30% of the observed variation. Individual gregariousness also had a small genetic component, with a low heritability estimate for degree (<5%). Centrality showed heritability up to 10% in one of three years sampled, whereas betweenness showed none, indicating modest genetic variation in individual sociability, but not group-switching tendencies. These findings suggest a small, albeit detectable, genetic influence on individual sociality, but pronounced spatial effects. Furthermore, our study highlights the importance of common environment effects (natal origin and brood identity), which essentially negated genetic effects when explicitly accounted for. In addition, we demonstrate that phenotypic resemblance can be a result of similarities beyond shared genes; spatial proximity at birth and natal environmental similarity explained up to 8% of individual sociability. Our results thus emphasise the role of non-genetic factors, particularly developmental and spatial variation, in shaping individual social behavioural tendencies.

## Introduction

Social behaviour is a pervasive aspect of animals’ lives, involving interactions with conspecifics through cooperation, mating, aggression, competition, and extending to activities like foraging in groups and anti-predator defence (Krause & Ruxton, 2002).The diversity of these interactions contributes to significant variation in the structure of animal societies. Understanding the causes and consequences of natural variation in group dynamics can provide valuable insights into the consequences of sociality, including effects on reproductive success, foraging efficiency and other selective benefits of grouping (Brown & Brown, 2000). For a comprehensive understanding of the evolution of sociality, however, it is crucial to explore the underlying causes of behavioural variation among and between individuals.

The social behaviour of an individual includes both its specific socio-behavioural characteristics and position within a social group, often called its social phenotype, along with its social environment, which is an amalgamation of social attributes and behavioural patterns exhibited by all individuals constituting a particular social group (Webber et al., 2023). The social environment serves as the contextual backdrop against which social phenotypes manifest and can thus constrain, as well as define, individual social attributes. Individuals will attempt to maximise their own fitness, and variable social phenotypes may be a consequence of individuals seeking optimal social niches through behavioural plasticity (Regan et al., 2022).This is when individuals adjust their behaviour both in response to and as a means of influencing varying environments, be it physical or social (Farine et al., 2015; Webber et al., 2023). What constitutes an optimal behavioural response may differ between individuals (Dingemanse & Wolf, 2013).

Individual-level variation in social behavioural phenotypes may show repeatability in its expression and can be influenced by genetic and environmental factors, or an interaction between the two. Within a population, individuals experience diverse environments, with small-scale variations in physical factors like habitat quality or weather conditions, or biotic influences like social interactions. These factors can create distinct phenotypes and contribute significantly to the observed variation (Cole et al., 2021; Regan et al., 2022). Environmental factors can even cause changes in an individual’s phenotype during growth and development (Dingemanse & Dochtermann, 2013). The rate of evolutionary response to selection depends on how heritable a trait is (Houle, 1992), which is why early research predominantly emphasised the genetic basis of traits. However, accurately describing how genotypes impact phenotypic variation necessitates acknowledging the role of environmental factors (Falconer, 1986). In this context, determining how repeatable/consistent a trait is, along with deciphering the relative contributions of genetic (e.g., additive genetic effects) versus non-genetic (e.g., resource distribution/developmental effects/phenotypic plasticity) sources of variation can help us understand the degree of behavioural plasticity exhibited by the individual, and set an upper limit to the heritability of the trait (Aplin et al., 2015).

In classical quantitative genetics, when comparing closely related individuals (which therefore share genes) to unrelated ones, similarities in phenotypes may imply a significant role for genes in shaping those traits, and the degree of familial resemblance in phenotype is given by the narrow sense heritability (h^2^), which is the proportional contribution of the additive genetic variance to the observed phenotypic variance (Wilson et al., 2010). Disentangling genetic versus non-genetic influences on behavioural traits can be challenging in wild animal populations. These challenges include generational overlap, unpredictable environmental factors, difficulties in trait measurement, limited pedigree data, complex population structures, covariation between genes and the environment, and the logistical difficulties associated with longitudinal studies. However, in populations with reliable parentage information, a mixed-effects modelling approach called the ‘animal model’ (Kruuk, 2004; Wilson et al., 2010) can be used. This model incorporates a relatedness matrix for all focal individuals and allows the inclusion of fixed and random effects that might be sources of variation for the trait, reducing bias in heritability estimates compared to other methods (Postma, 2014).

Furthermore, individuals may share similarities beyond shared genes, arising from experiencing similar small-scale environmental effects during development and adulthood. Behavioural variation within populations results from the interplay between permanent genetic, parental, developmental, or long-lasting environmental factors, alongside more flexible within-individual variation influenced by short-term environmental fluctuations or measurement error (Laskowski et al., 2022). Developmental noise arises from stochastic processes during an organism’s growth, leading to epigenetic modifications or variations in gene expression, resulting in different phenotypes among individuals with similar genotypes. These insights raise fundamental questions about the role of stochastic processes, particularly developmental variation, in shaping lifelong behavioural responses and trajectories.

While some studies have attempted to estimate the heritability of social-network derived social phenotypes (Brent et al., 2013; Godoy et al., 2022; Lea et al., 2010), this area remains relatively understudied due to the aforementioned constraints. In this study, we addressed this by leveraging a longterm monitored study system of wild great tits (*Parus major*) with detailed breeding records of individual birds since 1947 in Wytham Woods, Oxfordshire, UK (Lack, 1964). Along with a multigenerational pedigree, we used social network data obtained from a previous study on this system, which investigated the social behaviour of PIT-tagged individuals between the winters of 2011 to 2013 using RFID-antenna-fitted feeders, facilitating automated social data collection when individuals foraged together (Aplin et al., 2015).

Accordingly, this study had two primary objectives; (i) to estimate the repeatability (permanent environment effects) and narrow-sense heritability of group size choice and other social-network derived social phenotypes using a quantitative genetics animal model approach, and (ii) to decipher the role of non-genetic factors such as spatial effects, micro-environmental heterogeneity and developmental conditions in shaping individual social behavioural tendencies in the wild.

## Methods

### Study System

The analyses reported here use data from a long-term study on a population of great tits (*Parus major*) in Wytham Woods, Oxfordshire, U.K. As a cavity-nesting species, the majority of the breeding population uses nest boxes installed throughout the woods, which facilitates the trapping and monitoring of nestlings and adults during the breeding season (March-May each year), following a standardised protocol (Perrins, 1965). Captured birds are fitted with both a metal leg ring from the British Trust for Ornithology and, since 2007, with a plastic leg ring containing a uniquely identifiable passive integrated (PIT) tag from IB Technology, Aylesbury, U.K. Bird age and sex are determined upon capture using breeding records or plumage colouration. During winter, great tits form loose fission-fusion flocks of unrelated individuals, moving between patchy and ephemeral food sources, including bird feeders. To capture immigrant individuals (i.e., those not born and marked as nestlings in Wytham nestboxes), mist netting is performed at regular intervals, resulting in the ringing and PIT-tagging of a large majority of these birds.

### Social data

Data for this analysis were collected as part of a previous study (Aplin et al., 2015; Farine et al., 2015) on this system over three periods; 03 December 2011 – 26 February 2012, 01 December 2012 – 24 February 2013, and 30 November 2013 – 23 February 2014. This study spanned 13 weekends in each winter period (78 days of data collection, i.e., 26 days each year) where bird feeders fitted with radio-frequency identification (RFID) antenna, capable of scanning PIT tags, were deployed at 65 locations throughout the study site (Figure 1). Detailed spatiotemporal snapshots of individual foraging behaviours were recorded when individuals visited feeders and their unique PIT-tag code was scanned by the RFID antenna along with the associated time and location. A Gaussian mixture model was used to extract flocking events, which are high-density periods of feeding activity, from this spatiotemporal raw data through the *asnipe* package (Farine, 2013). Here, a cluster of visits to the feeder is identified as part of the same flocking event. This gave the group size for each flocking event, and there were repeated measures of it for each individual bird, corresponding with every flocking event it was part of. For a detailed description of this methodology, see Psorakis et al. (2012).

**Fig. 1.**
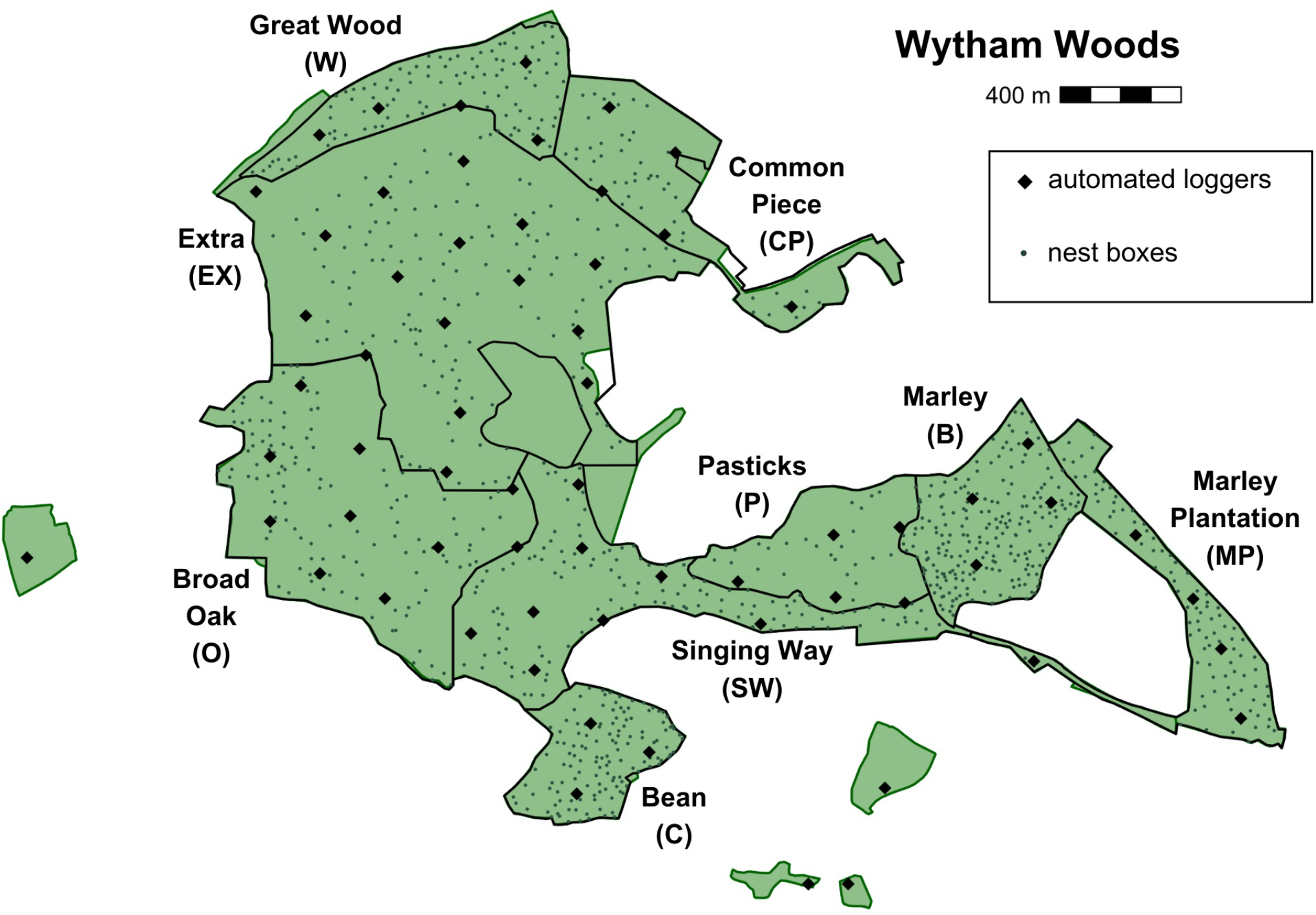
Map of the study site with the positions of 65 automated loggers, spaced approximately 250 m apart, which scanned PIT-tagged great tits over 13 weekends during 2011, 2012 and 2013 winters. Smaller points on the map indicate the locations of the 1020 artificial great tit-accessible nestboxes installed in the woodland, which is divided into 9 plots.

These flocking events provide a record of the individuals simultaneously or sequentially visiting the automated feeders, and it is assumed that birds that are part of the same flocking event at a feeder are associating with one another – in line with the ‘gambit of the group’ approach (Farine & Whitehead, 2015). Using this approach, social networks were created for each sampling period (each weekend), resulting in 39 total networks over three winters. Social network metrics were calculated for each individual that was part of those networks, which included the degree (an individual’s total number of associates), weighted degree (also called strength; a measure of number of associations weighted by their frequency), centrality (the eigenvector centrality; connectedness of an individual to other well-connected individuals), weighted centrality (how well connected an individual is to others with higher weighted degrees), and betweenness (the number of shortest paths that go through an individual). Additionally, flocking event data were pooled for each weekend to obtain an individual-level measure of average group size at weekend resolution over each winter.

While the degree and centrality reflect the overall gregariousness and sociability of an individual, betweenness represents an individual’s tendency to switch between groups. These metrics have been linked to several important processes, including the spread of social information (Aplin et al., 2015) and disease (Adelman et al., 2015; Ashby & Farine, 2022).

### Pedigree data

As part of the breeding survey, all individuals at nest boxes are identified either from their metal BTO rings on capture, or (in years since 2007) from PIT-tags detected by RFID antennas embedded in detachable ‘face-plates’ temporarily attached to a nest box. Birds without rings or tags are trapped and ringed along with their nestlings. Due to these efforts, a pedigree could be created using these data which assumes that the adult birds observed incubating or feeding chicks at a nest are the biological parents. Rates of extra-pair paternity in this population are relatively low (12.5%), and simulations suggest they do not significantly impact quantitative genetic methods or heritability estimates (Firth et al., 2015). The pedigree used for the analysis has 2825 total individuals (including only informative ones that contribute to subsequent analyses) and extends for up to 18 generations with 1552 maternities, 1476 paternities, 749 full siblings, 334 maternal half siblings, and 280 paternal half siblings. The focal individuals with social data have great-grandparents breeding from 1990 onwards, and thus, a fuller pedigree from 1980-2022 with 78,460 individuals was also used to compare the analyses between the pedigrees. However, the estimates remained unchanged and hence, the pruned pedigree with informative individuals was used for all the models reported here. 75% of the focal individuals have associated pedigree data. Of these, 51% were born in Wytham and thus have complete parentage information while the remaining 24% immigrated into the breeding population and have descendants in the pedigree. 25% of the focal birds have no associated pedigree data (i.e., they were observed feeding during winters but were neither offspring of other birds, nor contributed to the breeding population).

### Statistical Analyses

All analyses were conducted in R version 4.2.2 (R Core Team, 2022).

### Repeatability and Heritability of Social Phenotypes

ASReml-R software (Butler et al., 2007) was used to run animal models to partition the phenotypic variance into genetic and environmental variance components, in order to assess the repeatability and heritability of group size (Wilson et al., 2010). Separate models were run for the winters of 2011, 2012, and 2013 with 1,085 (343,458 observations), 720 (244,064 observations) and 789 (204,296 observations) individuals respectively to assess within-year variation in the social phenotype. Here, observation refers to each flocking event an individual was part of and thus, has a corresponding group size measure for. A model was also run for the entire dataset with year as a fixed effect, to assess between-year variation in group size for 1,823 individuals (791,818 observations, median; 356 observations per individual, IQR:468). Of the 1823 birds, 979 were seen in two out of three years, and 208 individuals were observed across all three winters.

Animal models were also run for six additional traits as response variables; namely average group size (group sizes from each flocking event averaged for each individual for every weekend they were observed in each winter), and five network traits obtained separately from the social networks created for 39 weekends across the three years. The entire dataset contained 1823 birds (23741 observations), and models were run separately for 2011, 2012 and 2013 with 1085 (9855 observations), 720 (6538 observations), and 789 (7348 observations) individuals respectively. Thus, social data was analysed at two levels; as repeated measures of group size for each individual every time it was part of a flocking event, and at the weekend-level through social networks generated from the same spatio-temporal flocking event data.

Repeatability quantifies the variation among repeated observations of the same individual which is reproducible, indicating the consistency of that individual phenotype (Nakagawa & Schielzeth, 2010). To estimate the repeatability of group size within and between years, individual identity was fitted as a random effect and the repeatability was calculated as the proportion of phenotypic variance (V_P_) explained by the individual variance (V_ID_). The repeatability or the intra-class correlation (ICC) was also estimated using the *rptR* package (Stoffel et al., 2017) to cross-check the estimates from ASReml-R.

To assess heritability, the pedigree was incorporated as a relatedness matrix linked to individual identity and fitted as a random effect to estimate the additive genetic variance (V_A_). The proportion of phenotypic variance explained by additive genetic effects is the narrow-sense heritability (h^2^) of the trait (V_A_/V_P_).

Furthermore, the feeder identity (logger) was fitted as a random effect to account for location-specific spatial effects and calculate the variation in group size that could be attributed to the location of the flocking events (V_LOG_).

Thus, the following models were run with the factors explained above for each winter (2011, 2012, 2013);

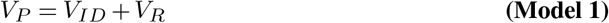

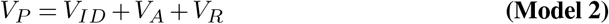

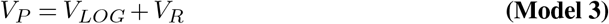

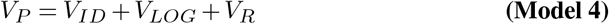

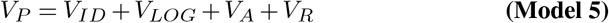

where V_R_ is the residual variance which accounts for the variation arising from environmental and other effects that have not been explicitly included in the model and V_ID_ is the permanent environment effect (individual identity) which is the non-additive contribution to fixed among-individual differences.

The aforementioned models were also run with the winter year included as a fixed effect, in order to control for the influence of year-to-year variation while assessing group size variance between-years. In doing so, the contribution of year of measurement to the variation in group size is essentially removed, isolating the effects of interest more accurately. For these models, the entire dataset across three years was used (with 1823 individuals) and the model structures were retained (Model 1–Model 5).

Using the same model framework, repeatability and heritability estimates were calculated for the weekend-level network traits as well. As before, the following models were run for the three winters separately and for the entire dataset with year as a fixed effect. In this case, as birds were observed over multiple locations, the V_LOG_ term was not fitted.

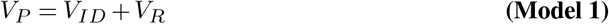

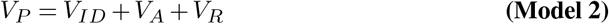

The proportion of variance explained by each of the variance components was calculated as the ratio of the relevant component to the total phenotypic variance (V_P_). To check for the significance of the random effects, likelihood ratio tests were performed by comparing models with and without the inclusion of the random effect term and comparing the log-likelihoods of the models while assuming a *χ*^2^ distribution with one degree of freedom (Wilson et al., 2010).

### Natal Effects on Social Behaviours

Of the 1823 individuals from the social network data, 938 birds (51.5%) were known to have been born in Wytham because they had been ringed as a nestling, and thus, had natal information available as part of the breeding survey. This includes the section of the woods each individual was born in (one of nine nestbox plots ranging from 18.7 to 120.6 hectares in size, which provide an approximation of broad-scale environmental heterogeneity within the study site; Figure 1), denoted here as ‘natal section’. Additionally, since each locally-born nestling is ringed when 14 days old, there is information available on the brood each individual belonged to (‘brood identity’). In the focal dataset, there are 558 unique broods having 1-7 focal individuals; 340 out of 938 birds have unique brood identities while the rest have 1-6 within-brood siblings within the social dataset.

ASReml-R software was used to model these two natal variables as random effects to estimate how much they contributed to variation in group size choice and degree (a measure of individual gregariousness) and to quantify the effect of common environments shared by individuals. Common environments, like the shared natal environment of siblings for instance, can increase phenotypic resemblance between relatives, and upwardly bias additive genetic variance (V_A_) unless explicitly accounted for (Kruuk & Hadfield, 2007). Group size choice and degree were selected for further analyses as they both showed considerably higher repeatability than the other traits (details in the results section).

The 938 locally-born birds were part of 401,915 flocking events across the three winters and had 12052 observations for degree via social network data at the weekend-level. As previously described, six models (Model a–Model f) were created for the variance decomposition of group size, with the winter year as a fixed effect.

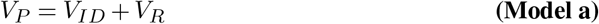

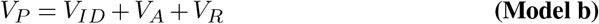

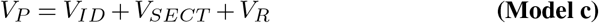

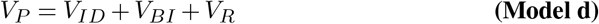

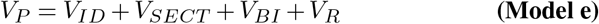

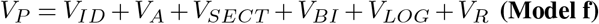

All six model structures were retained for degree as the response variable, apart from Model f from which the spatial location of loggers (V_LOG_) was removed.

For both traits, three additional models (Models g, h, i) were developed to model natal spatial information more directly. This was done by creating two types of matrices describing the similarity of non-genetic effects experienced by individuals during their development, specifically the fine-scale spatial effects (Jones et al., 2024).

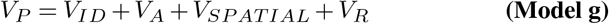

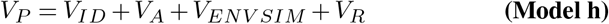

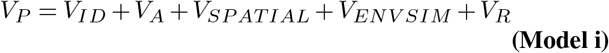

The first matrix, a spatial proximity matrix, was constructed by calculating the distance between the natal nestboxes of all possible pairs of individuals. These values were scaled and ranged from 0 to 1, where 0 represents the maximum distance between the natal nestboxes of two individuals and 1 represents the diagonal, indicating maximum spatial proximity with oneself.

The second matrix represented natal environmental similarity by combining five measures of small-scale environmental variation into an environmental similarity value between individuals. These measures included altitude, edge distance index, northness, oak-tree density within 75 meters, and population density (expressed as the square root of territory size). These factors have been shown to influence breeding behaviour in this great tit population and may also impact the developmental environment of chicks. Additionally, they are known to vary over small-scales in the woodland (Wilkin, Garant, et al., 2007; Wilkin, Perrins, & Sheldon, 2007; Wilkin et al., 2006). Each variable was standardized to have a mean of zero and a variance of one and the Euclidean distance between the environmental measures for each pair of individuals was then calculated. This distance represents the straight-line distance between two vectors of environmental measures in multivariate space. The resulting similarity matrix effectively captures the degree of environmental similarity experienced by individuals as a single value (see Thomson et al. (2018) for a description of this method).

Both these matrices can be incorporated as random effects in the animal model (similar to the genetic relatedness matrix) allowing us to estimate the contribution of natal environmental effects to overall phenotypic variance (Thomson et al., 2018). It would be fair to assume that birds born in proximity to one another would experience similar environments and thus, the inclusion of both these matrices might be seen as a redundant exercise. However, birds born spatially far apart can experience similar environments, and thus it is important to account for spatial and environmental similarities that extend beyond physical distance alone (Jones et al., 2024).

As previously described, log-likelihood ratio tests were performed on models with and without the random effects to check for the significance of the added terms.

## Results

### Repeatability and Heritability of Social Phenotypes

Within each winter, birds were significantly consistent in their group size with repeatabilities of 0.192, 0.303, and 0.205 for 2011, 2012 and 2013 respectively (Model 1). Between-year repeatability for individuals was 0.209. After incorporating pedigree information to explore additive genetic effects (Model 2), repeatabilities changed minimally but heritability estimates were small and close to zero (Figure 2 (a)).

**Fig. 2.**
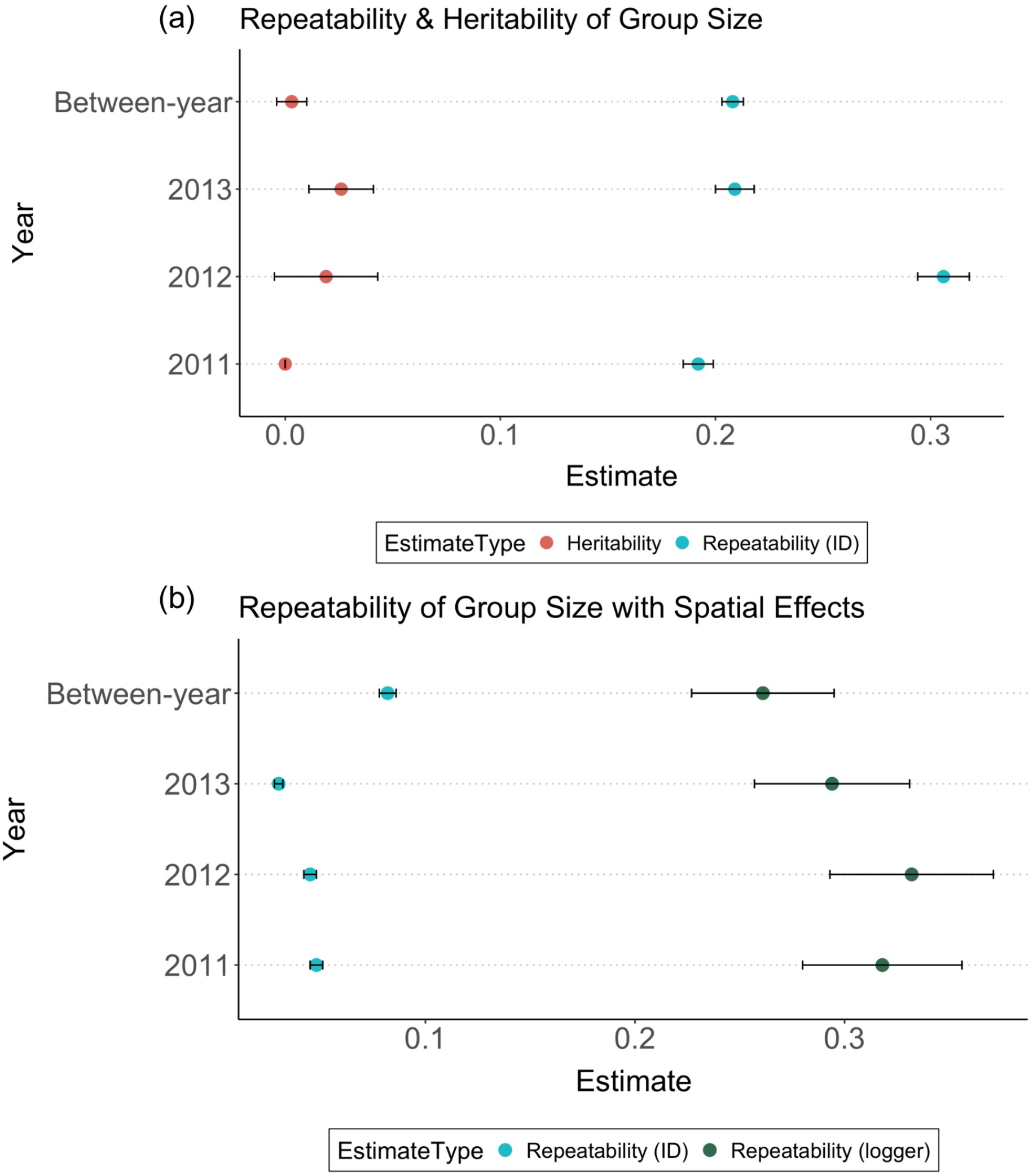
(a) Repeatability (shown in blue) and Heritability (shown in red) estimates for group size when random effects are individual identity (ID) and individual identity linked to a pedigree, showing within-year (2011, 2012 and 2013 winters) and between-year (across all 3 years) estimates (Model 2). (b) Repeatability estimates for individual identity (shown in blue) and spatial effects i.e., logger included as random effect (shown in green). Including logger as a random effect (model 4) greatly reduces estimates for individual-level repeatability. V_ID_= focal individual permanent environment effect, V_A_ = additive genetic effect, V_LOG_ = spatial effect of logger/feeder, V_R_= residual variance. Error bars show standard error.

Feeder (logger) identity was added as a random effect to explore variation in flocking behaviour at different locations. When a model was run with only logger as a random effect (Model 3), the repeatability of group size at loggers (i.e. with no individual identity data at all) was high for all years (0.245-0.296) which implies that there is consistency in the group sizes observed at a particular feeder and part of the variation can be attributed to the feeder the bird is feeding at. Consequently, both individual identity and logger were incorporated as random effects (Model 4) to estimate site-specific effects on group size variation. High repeatabilities were observed for loggers (feeder identity) while repeatabilities of group size for individuals reduced substantially, suggesting a large effect of location on the observed individual-level variation (Figure 2 (b)). Additionally, heritability estimates remained virtually zero even when site-specific effects were accounted for (Model 5).

The breakdown of phenotypic variance in group-size variation into its various components revealed that the contribution of additive genetic variance (V_A_) is extremely small, while variances attributed to individual identity and logger identity are larger (Figure 3, Table 1, supplementary information). In this case, for the group size phenotype which is repeatable at the individual level, genetic factors have minimal influence and the variation is largely explained by the effect of the environment.

**Fig. 3.**
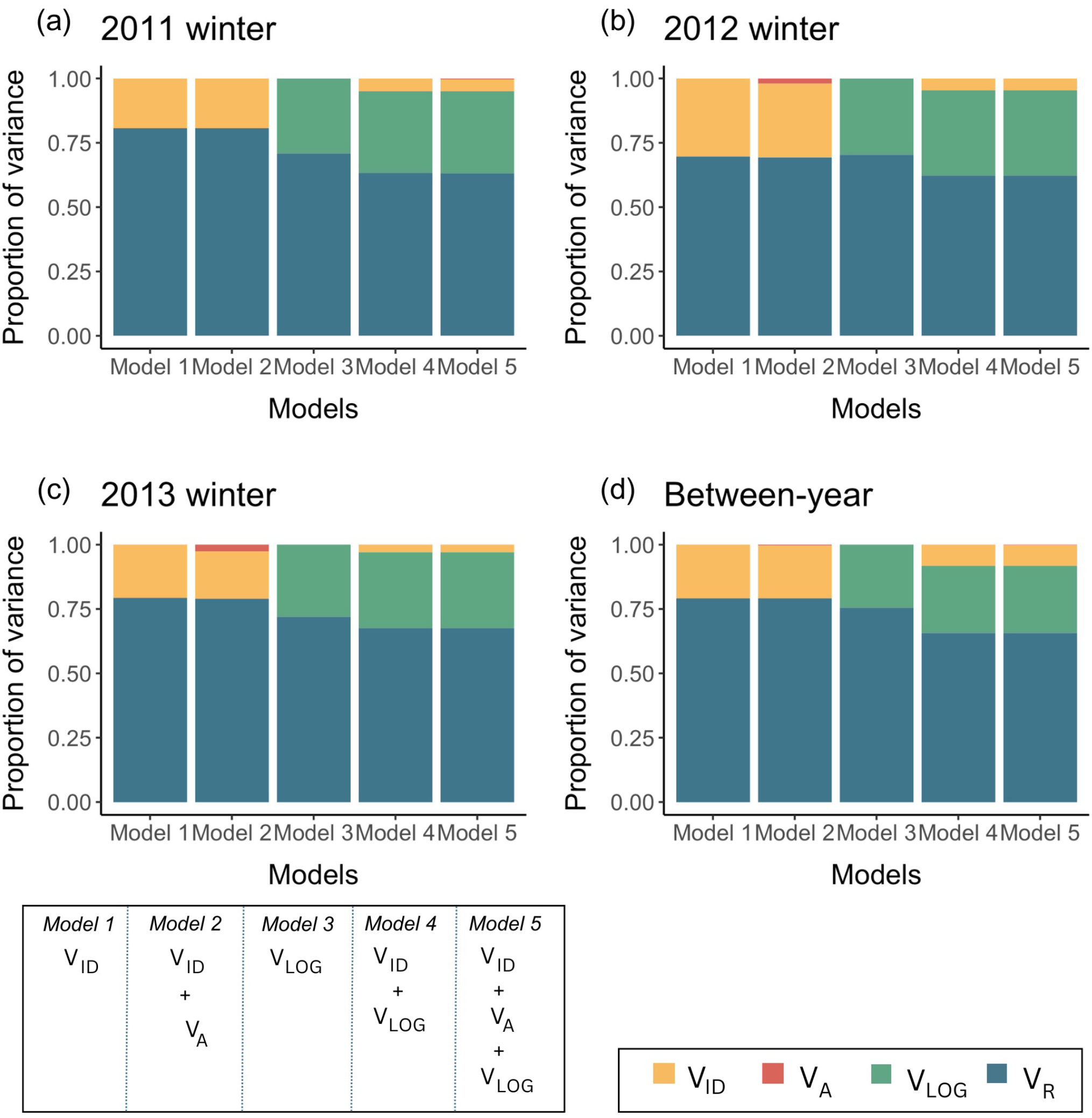
Proportion of variance components from the 5 different models for group size in (a) 2011–2012 winter; (b) 2012–2013 winter; (c) 2013–2014 winter; (d) across 3 years (between-year repeatability). V_ID_ = focal individual permanent environment effect, V_A_ = additive genetic effect, V_LOG_ = spatial effect of logger/feeder, V_R_ = residual variance. Specific information for model structures is available in the methods.

Using the same model framework, when data were pooled for each weekend to calculate the average group size at this level, repeatability estimates were 0.379, 0.586, and 0.545 for the three years (Figure 4, and Table 2, supplementary information). These estimates are quite comparable to the previous study that estimated the repeatability of average group size in this system, which also accounted for local population size and network density (0.43, 0.64 and 0.60 for 2011, 2013 and 2013 respectively (Aplin et al., 2015)); the larger estimates are expected as averaging across weekends reduces the sampling variance. A similar trend was observed for degree; 0.379, 0.574 and 0.493 respectively (Table 2, supplementary information), and comparable to the estimates from the earlier study (0.46, 0.61, and 0.58).

**Fig. 4.**
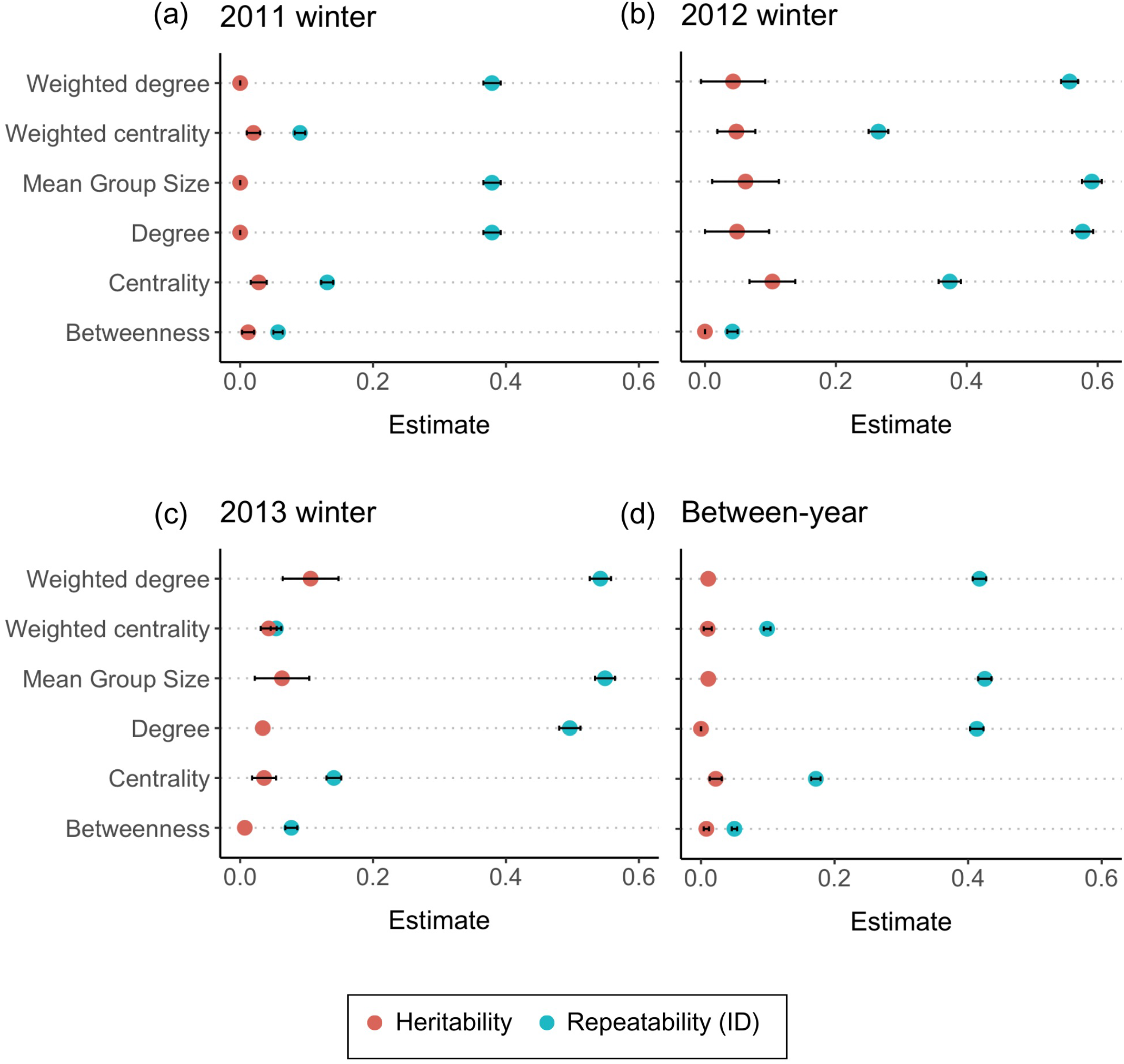
Repeatability (in blue) and narrow-sense heritability (in red) estimates for social network traits in (a) 2011–2012 winter; (b) 2012–2013 winter; (c) 2013–2014 winter; (d) across 3 years (between-year repeatability) using the model; V_P_ = V_ID_ + V_A_ + V_R_. V_ID_ = focal individual permanent environment effect, V_A_ = additive genetic effect, V_R_ = residual variance, V_P_ = total phenotypic variance. Repeatability of the social network trait for focal individuals is calculated as proportion of V_ID_ +V_A_ to V_P_, h^2^ = narrow-sense heritability as proportion of V_A_ to V_P_.

Mean group size, degree and weighted degree showed high repeatability (0.3-0.5), but have low heritability values (<0.06); the only exception being a heritability estimate of 11% for weighted degree in the winter of 2013. (Figure 4 (c)). Centrality and weighted centrality, on the other hand, showed lower repeatability (0.1-0.3) but heritability values ranging from 0.01-0.103, signifying a modest, but detectable genetic component of variance between years. In contrast, betweenness showed very little consistency across all sampling periods, with repeatability estimates <0.076 and virtually zero heritability (Table 2, supplementary information).

### Natal Effects on Social Behaviours

Group size choice and degree have moderate repeatability between years, with individual identity explaining 20.6 ± 0.8% and 41.5 ± 1.4% of the phenotypic variation respectively (Supplementary Table 1 and 2). These are estimates for when only the permanent environment effect of individual identity is considered, and they remained consistent when a subset of locally-born birds was used for the analyses (Model a). However, restricting the analyses to the 938 Wytham-born individuals increased heritability estimates slightly from 0.003 to 0.031 for group size and from 5.89 × 10-8 to 0.041 for degree (Model b), when compared with the full dataset (1023 birds). When natal environmental effects were incorporated in the models, effectively all of the additive genetic variance was replaced by variation attributable to natal section and brood identity (Model e), as depicted in Figure 5.

**Fig. 5.**
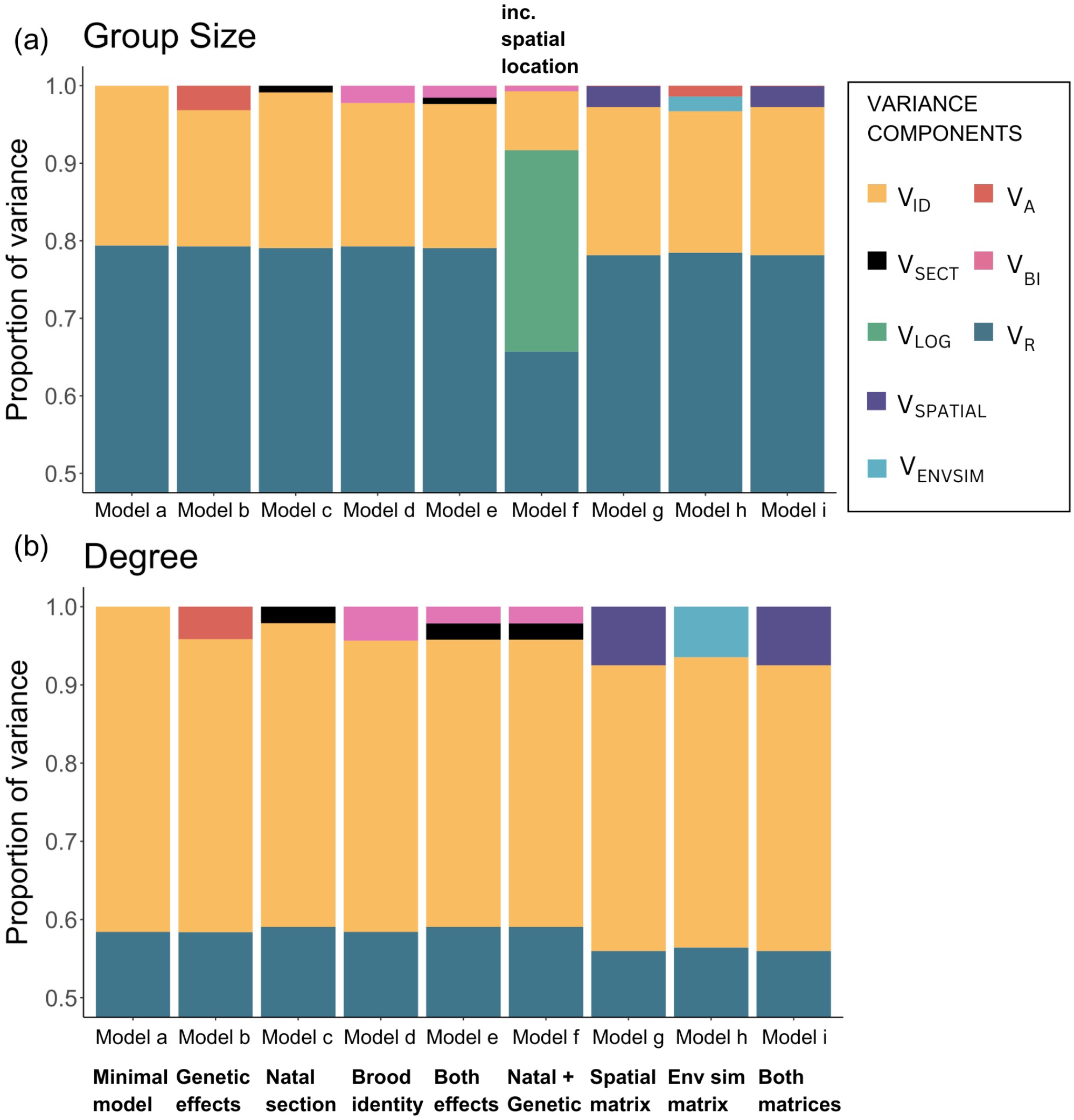
Proportion of variance components from the 9 different models for group size and degree across 3 years (winters). All models have year as a fixed effect and follow the same model structures (detailed in the methods) for both the phenotypes, with the exception of an additional term for spatial location of loggers in Model f for group size. V_ID_ = focal individual permanent environment effect, V_A_ = additive genetic effect, V_SECT_ = natal section, V_BI_ = brood identity, V_LOG_ = spatial effect of logger/feeder, V_SPATIAL_ = spatial proximity matrix, V_ENVSIM_ = natal environmental similarity matrix, V_R_ = residual variance.

In the group size models, addition of the natal section effect (V_SECT_) (Model c) to the minimal model (Model a), explained 0.8% of the group size variation, minimally reducing the permanent environment effect (V_ID_) from 20.6 to 20%. Brood identity (V_BI_), however, explained 2.2% of the variation, further decreasing V_ID_. When both natal effects were included in the model, they cumulatively explained 2.3% of the variation and could be explaining some of the additive genetic variance in Model b (3.1%). As expected, adding the natal and genetic terms into the model reduced the heritability estimate to virtually zero, indicating that almost all of the V_A_ in Model b was upwardly biased due to lack of accounting for common environment effects. As previously shown, incorporating the spatial effect, V_LOG_ here (Model f) considerably reduced the permanent environment effect, V_ID_ (from approximately 20% to 7.6%).

The influence of natal effects on degree was broadly similar. The natal section (V_SECT_) and brood identity (V_BI_) individually explained 2.1 ± 1.3% and 4.3 ± 2.3% respectively (Model c and Model d) and addition of these random terms significantly improved the models. Of the 41.5% attributed to individual effects (V_ID_) (Model a) 4.1% was explained by additive genetic variance (Model b) which reduced to 3.2 × 10-8 when natal environmental effects were jointly incorporated (Model f). As before, the additive genetic variance was much higher when natal effects were not modelled and this highlights the importance of accounting for sources of variation that could bias heritability estimates.

Heritability estimates remained negligible when the spatial proximity and natal environmental similarity matrices were included for both group size choice and degree. In fact, the matrices explained substantially more of the phenotypic variance than the cumulative effects of both natal section and brood identity. The spatial proximity between natal nestboxes (V_SPAT_) explained 7.4 ± 3.9% and 2.7 ± 1.6% of the total variance in degree and group size respectively (Model g). The natal environmental similarity matrix (V_ENVSIM_) explained slightly less of the overall variance than the other matrix but still had a notable contribution. It explained 6.4 ± 3.3% and 1.9 ± 1.3% of the variation in degree and group size choice respectively (Model h). Not only do these matrices account for the common environment effect (which natal section and brood identity were a part of), they also explain some of the residual variation for both these traits.

Subsequently, the final multi-matrix model included both the spatial proximity and natal environmental similarity matrix along with the genetic relatedness matrix (Model i). Here, heritability estimates remained close to zero and the inclusion of the spatial proximity matrix markedly reduced the contribution of the natal environmental similarity matrix to practically zero in both traits. The spatial proximity matrix explained 7.4% of the variation in degree and 2.7% of the group size variation, indicating that a substantial portion of the phenotypic variance can be explained by the spatial location of where individuals were born and the common environments shared by not just siblings, but also by birds that were born in close proximity to one another. Figure 5 represents how all these natal effects on adult phenotypes can be disentangled.

## Discussion

Consistent with previous studies on this system, individual-level characteristics significantly impacted group size and shaped the formation of foraging flocks in great tits. Individual great tits were substantially repeatable in their group size choice (R = 0.19 – 0.3) and other social-network traits, namely degree (R = 0.3 – 0.5) and centrality (0.1 – 0.3); these figures are broadly in line with a meta-analysis across several taxa that estimated an average repeatability of 0.37 for all behaviour (Bell et al., 2009). Disentangling the genetic and non-genetic sources of variation in these behaviours is challenging but crucial to improve our understanding of the formation and maintenance of social structures in individuals’ social networks.

By combining a multi-generational pedigree and a large observational dataset for social behaviours in wild great tits, we used animal models to partition the observed variation in group size into genetic and environmental sources. This analysis demonstrated very low heritability for group size choice in wild great tits, with additive genetic variance accounting for less than 2% of the total phenotypic variance on both short and long-term timescales. This trend continued with social network traits also having minimal heritability values. Degree and weighted degree had small genetic components (<5%) with the exception of weighted degree having a heritability estimate of 10.6% in the winter of 2013. Heritability values also showed variation between years for centrality and weighted centrality, ranging from 0 to 10.3%. However, betweenness showed very low repeatability and heritability across all years. These findings indicate a small genetic basis for the quality, quantity and strength of connections individuals have but not for their tendencies to switch between social groups.

Other studies that have attempted to assess genetic variation in social phenotypes derived from social networks have shown considerable heritability for some metrics. Affiliative and aggressive social behaviours in rhesus macaques (*Macaca mulatta*) (Brent et al., 2013) have heritability estimates of 0.84 and 0.66 respectively, yellow bellied marmots (*Marmota flaviventris*) (Lea et al., 2010) have estimates around 0.11 and a recent study on white-faced capuchin monkeys (*Cebus capucinus imitator*) (Godoy et al., 2022) placed the heritability of sociality at 0.152. While being higher than the estimates from this study on great tits, it is important to consider that social groups in marmots, macaques and capuchins have strong relatedness in their composition, and hence, heritability estimates can be influenced by kin-biases despite these studies having partly corrected for individuals sharing physical space within a group (Radersma, 2021).

Group size is often quite flexible in many species (Krause & Ruxton, 2002); for instance, shoaling fish are known to change group size in response to manipulated environmental conditions (Hoare & Krause, 2003). Despite this adaptability, groups generally form as individuals strive to enhance their own fitness while balancing the costs and benefits associated with grouping, implying that social structures emerge when individuals adjust their behaviour dynamically to fit their preferred group size, which may change depending on environmental shifts, group compositions or trait differences among individuals within groups (Couzin & Krause, 2003). Wytham Woods is heterogeneous over small scales, and the woodland exhibits variations in microhabitat quality and local population density. As a result, the consistency of flocking behaviour may be influenced by individual differences in space use (Shizuka et al., 2014). Approximately 30% of the variation in group size could be attributed to the spatial location of the flocking event. As is apparent from Figure 3, spatial effects seem to explain a considerable amount of individual-level variation while also explaining some of the residual variation that can be attributed to environmental effects.

Birds belonging to foraging flocks access ephemeral food sources as they move across the environment while also sharing the responsibility of looking out for predators. While these group behaviours can be analysed broadly, social network analysis allows us to look at fine-scale non-random individual-level interactions at a spatio-temporal level (Farine & Whitehead, 2015). However, since social network analysis interprets associations among individuals based on their spatial proximity, a key challenge is distinguishing whether these associations are because of shared spatial preferences, favoured social associations, or both (Webber et al., 2023). Repeatability at individual loggers is high, indicating that specific locations show consistency in group sizes. Accounting for spatial environments and other factors is important because individuals in high-density populations are more likely to be socially connected than those in lower-density populations. The density of individuals at the feeder is a condition in itself, as it changes the risk perception, resource availability, and the social environment of the spatial location (Webber et al., 2023). Combined with other factors like habitat, time of day/year, information availability and other environmental constraints, spatial effects on group size variation could be a result of the conditions at the foraging location itself or individual decision-making.

Tits use social information while making decisions about dispersal and settlement, with individual differences in information use. Additionally, they use social information to discover food patches through local enhancement and social learning (Aplin et al., 2012). Consequently, a component of an individual’s social phenotype may be due to its choice of when and where to forage and individual settling decisions (Aplin et al., 2015). Moreover, if animals vary in their degree of sociality, differences in association strengths could influence group-level movements, with repeatable individual-level leader-follower characteristics impacting group dynamics. While the foraging location seems to have a notable impact on the consistency of an individual’s group size, additional factors beyond spatial constraints may play a significant role in explaining the observed variation in group size, including individual characteristics, social interactions, environmental cues, climatic conditions, natal effects, or other unaccounted for local aspects.

Including natal factors into the models partly corroborated this idea; the birth location of an individual, which is broadly indicative of the habitat experienced by a bird during development, explained 3-5% of the repeatability in group size choice and degree. Moreover, up to 10% of the permanent environment of the individual could be explained by the identity of the brood it belonged to. Thus, individual-level characteristics contributing to consistent social phenotypes seem to largely be shaped by the physical and social environment during development. Furthermore, accounting for these natal effects completely negated the influence of any significant genetic effects, emphasising the importance of considering shared environments between related individuals. In natural populations, individuals may have similar phenotypes not just because they share genes, but also because they share common environments, such as a parent or a nest for instance. This can complicate the distinction between genetic and environmental influences, and it is essential to account for these common environment effects to prevent obtaining inflated heritability estimates.

Our findings further demonstrated the importance of accounting for fine-scale spatial variation in shaping permanent differences between individuals, which are often overlooked in traditional animal models (Stopher et al., 2012). Common environment effects can manifest beyond spatial proximity between individuals, if they share similar microenvironments despite being far apart in space. Natural environments are often heterogenous across spatial scales that are relevant for their inhabitants. This may lead to a scenario where individuals that are close in space experience quite different micro-environments (Jones et al., 2024). Through this study, we explicitly modelled these fine-scale environmental effects experienced by individuals during their development by using spatial proximity and natal environmental similarity matrices to show that natural variation in early-life environments can be vastly indicative of individuals’ social trajectories.

It is apparent from this study that such non-genetic sources of variation might be the key drivers of variation in social behaviours, with genetic influences playing a minimal role. A critical limitation to address in this analysis would be to identify and account for indirect genetic effects. Social network metrics are a consequence of individual behaviour in the context of other interacting individuals, each with their own phenotype. Heritability estimates would be more accurate if the additive genetic variation in the phenotypes of the interacting individuals is also accounted for (Godoy et al., 2022).

Overall, our findings highlight the complex interplay between individual characteristics, spatial location, and other contextual factors in shaping consistency in social behavioural traits. Future research could delve deeper into understanding these underlying mechanisms and identifying the specific factors driving within-individual consistency in social behaviour within local contexts. Furthermore, our study emphasises the need for incorporating detailed environmental effects into models which, albeit challenging, is crucial for reducing bias and improving the accuracy of quantitative genetic analyses.

## ACKNOWLEDGEMENTS

We are grateful to the many people who helped collect these data over the years: numerous people who collected long-term data from the Wytham tit study which supported the pedigree construction, and members of the EGI Social Network group who collected the social network data, and particularly Damien Farine and Josh Firth for work on estimating social phenotypes. The long-term population study has been supported by numerous funding sources, including recently by grants from BBSRC (BB/L006081/1) and NERC (NE/K006274/1, NE/S010335/1); the social network data were funded by an ERC grant (AdG250164). We would also like to thank Carys Jones for providing some of the code used for analyses and for useful inputs. Devi Satarkar is supported by the Oxford-Oxitec Graduate Scholarship. Irem Sepil is supported by a Royal Society Dorothy Hodgkin Fellowship (DHF111084).

## Author contributions

All authors contributed to the ideas of this study. DS conducted the statistical analyses with critical inputs on methodology from BCS. DS drafted the manuscript with IS and BCS providing substantial feedback.

## Data and code availability

Data and code to reproduce all analyses are available at https://github.com/devisatarkar/SocNet_ASRemL_herit-natal

