## Supplementary Information for "Genetic, Natal, and Spatial Drivers of Social Phenotypes in Wild Great Tits"

**Table 1.** Variance components from 5 animal models (with various combinations of random effects used) detailing the estimates (standard error). V_ID_ = focal individual permanent environment effect, V_A_ = additive genetic effect, V_LOG_ = spatial effect of logger/feeder, V_R_ = residual variance, V_P_ = total phenotypic variance (sum of all variance components), R_ID_ = Repeatability of group size for focal individuals given as proportion of V_ID_ or V_ID_ +V_A_ to V_P_, R_LOG_= Repeatability of group size at loggers given as proportion of V_LOG_ to V_P_, h^2^ = narrow-sense heritability as proportion of V_A_ to V_P_.

| **Model** | **Year** | **2011** | **2012** | **2013** | **All 3 (FE=year)** |
| --- | --- | --- | --- | --- | --- |
| Random effect = ID  (*Model 1)* | V_ID_ | 4.548 (0.206) | 5.999 (0.328) | 2.891 (0.157) | 4.205 (0.147) |
|  | V_R_ | 19.028 (0.045) | 13.739 (0.039) | 11.153 (0.034) | 15.99 (0.025) |
|  | V_P_ | 23.576 | 19.738 | 14.044 | 20.195 |
|  | R_ID_ | **0.192** (0.007) | **0.303** (0.011) | **0.205** (0.008) | **0.209** (0.005) |
| Random effect = ID + pedigree  (*Model 2*) | V_ID_ | 4.548 (0.206) | 5.673 (0.498) | 2.589 (0.221) | 4.159 (0.187) |
|  | V_A_ | 5.948 x 10^-7^ | 0.392 (0.489) | 0.366 (0.225) | 0.06 (0.153) |
|  | V_R_ | 19.028 (0.045) | 13.739 (0.039) | 11.153 (0.034) | 15.99 (0.025) |
|  | V_P_ | 23.576 | 19.804 | 14.108 | 20.209 |
|  | R_ID_ | 0.192 (0.007) | 0.306 (0.012) | 0.209 (0.009) | 0.208 (0.005) |
|  | h^2^ | 2.523 x 10^-8^ (2.26 x 10^-10^) | 0.019 (0.024) | 0.026 (0.015) | 0.003 (0.007) |
| Random effect = logger  (*Model 3*) | V_LOG_ | 7.230 (1.280) | 4.920 (0.851) | 3.842 (0.682) | 4.899 (0.866) |
|  | V_R_ | 17.523 (0.042) | 11.718 (0.033) | 9.901 (0.03) | 15.094 (0.023) |
|  | V_P_ | 24.753 | 16.638 | 13.743 | 19.993 |
|  | R_LOG_ | 0.292 (0.036) | 0.296 (0.036) | 0.279 (0.035) | 0.245 (0.032) |
| Random effect = ID + logger  (*Model 4*) | V_ID_ | 1.300 (0.064) | 0.813 (0.049) | 0.428 (0.027) | 1.767 (0.065) |
|  | V_LOG_ | 8.442 (1.498) | 5.996 (1.06) | 4.191 (0.746) | 5.624 (0.995) |
|  | V_R_ | 16.804 (0.040) | 11.235 (0.032) | 9.604 (0.03) | 14.13 (0.022) |
|  | V_P_ | 26.546 | 18.044 | 14.223 | 21.521 |
|  | R_ID_ | 0.048 (0.003) | 0.045 (0.003) | 0.030 (0.002) | 0.082 (0.004) |
|  | R_LOG_ | 0.318 (0.038) | 0.332 (0.039) | 0.294 (0.037) | 0.261 (0.034) |
| Random effect = ID + logger + pedigree  (*Model 5*) | V_ID_ | 1.209 (0.086) | 0.813 (0.049) | 0.428 (0.027) | 1.735 (0.083) |
|  | V_LOG_ | 8.445 (1.498) | 5.996 (1.064) | 4.191 (0.746) | 5.624 (0.995) |
|  | V_A_ | 0.123 (0.086) | 2.2 x 10^-7^ | 1.5 x 10^-7^ | 0.041 (0.070) |
|  | V_R_ | 16.804 (0.040) | 11.233 (0.032) | 9.604 (0.03) | 14.130 (0.022) |
|  | V_P_ | 26.581 | 18.042 | 14.223 | 21.53 |
|  | R_ID_ | 0.050 (0.003) | 0.045 (0.003) | 0.030 (0.002) | 0.082 (0.004) |
|  | R_LOG_ | 0.317 (0.038) | 0.332 (0.038) | 0.294 (0.037) | 0.261 (0.034) |
|  | h^2^ | 0.004 (0.003) | 1.22 x 10^-8^ (6.9 x 10^-10^) | 1.06 x 10^-8^ (5.5 x 10^-10^) | 0.001 (0.003) |

**Table 2.** Repeatability & narrow-sense heritability estimates for social network traits within-years (2011, 2012, and 2013) and between-year using two models; V_p_ = V_ID_ + V_r_ shown as V_ID_, and V_p_ = V_ID_ + V_A_ + V_r_ shown as V_ID_ + V_A_. V_ID_ = focal individual permanent environment effect, V_A_ = additive genetic effect, V_R_ = residual variance, V_P_ = total phenotypic variance, R_ID_ = Repeatability of group size for focal individuals given as proportion of V_ID_ or V_ID_ +V_A_ to V_P_, h^2^ = narrow-sense heritability as proportion of V_A_ to V_P_.

| **Trait** | **Model** | **Estimate** | **2011** | **2012** | **2013** | **Between-year** |
| --- | --- | --- | --- | --- | --- | --- |
| Mean group size | V_ID_ | R | 0.379 (0.013) | 0.586 (0.015) | 0.545 (0.015) | 0.423 (0.01) |
|  | V_ID_ + V_A_ | R | 0.379 (0.013) | 0.591 (0.015) | 0.549 (0.015) | 0.425 (0.01) |
|  |  | h^2^ | 4.42 x 10^-8^ (9.65 x 10^-10^) | 0.062 (0.051) | 0.063 (0.041) | 0.0109 (0.017) |
| Degree | V_ID_ | R | 0.379 (0.013) | 0.574 (0.015) | 0.493 (0.015) | 0.413 (0.01) |
|  | V_ID_ + V_A_ | R | 0.379 (0.013) | 0.577 (0.016) | 0.496 (0.016) | 0.413 (0.01) |
|  |  | h^2^ | 5.3 x 10^-8^ (1.16 x 10^-9^) | 0.049 (0.049) | 0.034 (0.036) | 5.89 x 10^-8^ (10^-9^) |
| Weighted Degree | V_ID_ | R | 0.375 (0.013) | 0.554 (0.0157) | 0.534 (0.0151) | 0.416 (0.01) |
|  | V_ID_ + V_A_ | R | 0.379 (0.013) | 0.557 (0.0136) | 0.542 (0.016) | 0.417 (0.010) |
|  |  | h^2^ | 5.1 x 10^-8^ (1.11 x 10^-9^) | 0.043 (0.049) | 0.106 (0.042) | 0.0108 (0.016) |
| Weighted Centrality | V_ID_ | R | 0.086 (0.007) | 0.259 (0.014) | 0.046 (0.007) | 0.097 (0.005) |
|  | V_ID_ + V_A_ | R | 0.09  (0.008) | 0.265 (0.015) | 0.054 (0.008) | 0.099 (0.005) |
|  |  | h^2^ | 0.02  (0.01) | 0.048 (0.029) | 0.043 (0.012) | 0.010 (0.006) |
| Centrality | V_ID_ | R | 0.126 (0.009) | 0.365 (0.016) | 0.136 (0.01) | 0.169 (0.007) |
|  | V_ID_ + V_A_ | R | 0.131 (0.009) | 0.374 (0.017) | 0.141 (0.011) | 0.172 (0.007) |
|  |  | h^2^ | 0.028 (0.012) | 0.103 (0.035) | 0.036 (0.018) | 0.022 (0.009) |
| Between-ness | V_ID_ | R | 0.0551 (0.006) | 0.0423 (0.008) | 0.076 (0.008) | 0.049 (0.004) |
|  | V_ID_ + V_A_ | R | 0.057 (0.007) | 0.042 (0.008) | 0.077 (0.009) | 0.050 (0.004) |
|  |  | h^2^ | 0.0123 (0.009) | 1.6 x 10^-8^ (2.97 x 10^-10^) | 0.007 (0.012) | 0.008 (0.004) |
